# Do Walking Muscle Synergies Influence Propensity of Severe Slipping?

**DOI:** 10.1101/776732

**Authors:** Mohammad Moein Nazifi, Kurt Beschorner, Pilwon Hur

## Abstract

Slipping is frequently responsible for falling injuries. Preventing slips, and more importantly severe slips, is of importance in fall prevention. Our previous study characterized mild slipping and severe slipping by the analysis of muscle synergies. Significant discrepancies in motor control of slipping have been observed between mild and severe slippers. We are further interested in whether differences exist in baseline motor control patterns between persons who experience mild and severe slips when exposed to a slippery contaminant. This study investigated walking with a muscle synergy approach to detect if walking muscle synergies differ between groups experiencing different slip severities. Twenty healthy young adults (8 mild slippers and 12 severe slippers) participated in this study and their muscle synergies of walking were extracted. Muscle synergy analysis showed that mild slippers had a higher contribution of hamstring and quadriceps during walking while severe slippers had increased contribution of tibialis group. This study provides novel information that may contribute to identifying diagnostic techniques for identifying persons or populations with a high risk of fall based on their walking patterns.

## 1 Introduction

In 2015, 17% of the fatal injuries were caused by slips, trips, and falls (Bureau of Labor Statistics US Department of Labor 2016a). Also, 27% of the total non-fatal occupational injuries in 2015 were caused by slips, trips, and falls, emphasizing the adverse consequences of the slipping and falling. Most importantly, “fall on the same level” was identified as the leading event contributing to days-away-from-work cases during 2015 (Bureau of Labor Statistics US Department of Labor 2016b). Slips are the main trigger to falling and considering the detrimental effect of the falls, confronting slips, as the main contributors to falling, should be a priority. Additionally, not all slips impose the same risk. “Severe slips” are more prone to result in a fall compared to “mild slips”. Hence, studying “severe slips” (in order to prevent them) is a necessary step toward fall prevention.

Numerous research articles have argued that motor control of the body’s complex muscular system is reduced to a lower dimensional modular control by the Central Nervous System (CNS) (Ting and Macpherson 2005; Tresch and Jarc 2009; d’Avella, Saltiel, and Bizzi 2003). These modules, also known as *muscle synergies*, have been extracted for various motor tasks (e.g. walking, slipping, swimming (Clark et al. 2010; d’Avella and Bizzi 2005;Nazifi et al. 2017;Nazifi, Beschorner, and Hur 2017)), and can potentially represent physical sub-tasks of the original motor task (Neptune, Clark, and Kautz 2009;Nazifi et al. 2017). On the other hand, prior research has indicated the potential influence of motor control during gait on an individual’s risk of falling. Moyer et al. (2006) used kinematic metrics of human gait (e.g. cadence) to evaluate an individual’s risk of experiencing a severe slip, indicating the link between gait parameters and slip severity. Given that muscle activations can imperatively affect the resulting kinematics, one may suspect that a similar link might relate slip severity to the lower extremity muscle activation patterns during walking. This speculation is also substantiated by our previous study claiming that the CNS uses the same patterns/modules to control both human walking and slipping based on a muscle synergy approach (Nazifi et al. 2017). Although our previous studies extracted walking muscle synergies, it is still unknown if the walking muscle synergies differ for individuals with different slip severity. Such knowledge is valuable as it may potentially result in a novel diagnosis method that only relies on walking behavior of subjects to eventually predict their slip severity.

In sum, this study intends to understand how muscle synergies observed during *<u>walking</u>* differ for the individuals who were classified as *<u>severe slippers</u>* compared to those who were classified as *<u>mild slippers</u>*. We hypothesize that muscle synergies of walking will differ between mild and severe slippers. Since the walking muscle synergies represent the neural control of the gait, the observed differences in different severities may potentially show the effect of the neural control of the gait on slip severity. Such knowledge is valuable since each muscle synergy is shown to be associated with a physical sub-task of a gross motor task (Nazifi et al. 2017). Hence, comparing walking muscle synergies is equivalent to identifying the walking subtasks that differ between mild and sever slippers. These differences in walking muscle synergies may help pinpoint the underlying limb coordination and walking habits that may contribute to a higher risk of fall on the slippery surface and can potentially be used in the development of programs for slip/fall prevention, diagnosis, and rehabilitation.

## 2 Materials and Methods

### 2.1 Subjects

A total number of twenty healthy young adults (9 females, 11 males) with an average age of 23.6 years old (SD = 2.52 years) participated in this study. Our previous studies (Nazifi, Beschorner, and Hur 2017;Nazifi et al. 2017) showed that 18 subjects can provide enough power (effect size = 1.28, power = 0.8) for statistical analysis. Subjects had no history of illnesses affecting gait (e.g., musculoskeletal, neurological, cardiovascular). All subjects signed the written consent forms prior to participation in this IRB-approved experiment at the University of Pittsburgh. Upon a secondary approval from IRBs of University of Pittsburgh and Texas A&M University, the anonymized data were analyzed in Texas A&M University for the current study.

### 2.2 Procedures

Subjects were asked to walk at their comfortable speed along a ten-meter pathway with an embedded force plate at the middle. There were two or three practice trials before the main walking trial (i.e. data recording trial). The starting point was adjusted in each trial to make subjects step on the force plate with their right limb. All subjects were provided with Polyvinyl chloride (PVC) soled shoes in their size to control the coefficient of friction across all the subjects. Subjects donned a harness to protect them from any potential injuries due to slipping during testing.

After the practice trial, subjects performed a normal walking trial during which the EMG signals and markers data were recorded for synergy extraction. After the normal walking trial, without informing the subjects of a change in walkway condition, a slippery solution was applied to the force plate. The slippery solution was a diluted glycerol solution, with 75% glycerol and 25% water, that has shown promise in providing a slippery surface by other researchers (Moyer et al. 2006; Chambers and Cham 2007; Beschorner and Cham 2008). The coefficient of friction was 0.53 and 0.03 for the dry and slippery conditions, respectively. Then, subjects performed an unexpected “slip trial” to classify the subjects into different severity groups (Figure 1). To minimize the audible and visible cues and ensure an unexpected slip, we administered the following: we dimmed the lights during the whole experiment. Also, between the trials, subjects were asked to look away from the walkway, while listening to loud music for one minute. Next, subjects were asked to turn around to face the walkway again, place their feet on the instructed location, and start walking on the signal. Subjects were also asked to keep their eyes at an eye-level target on the wall (i.e. horizon). The slipping trial was only recorded to classify subjects into either mild or severe slippers while the main data analyzed in this study was the walking behavior of the subjects.

**Figure 1:**
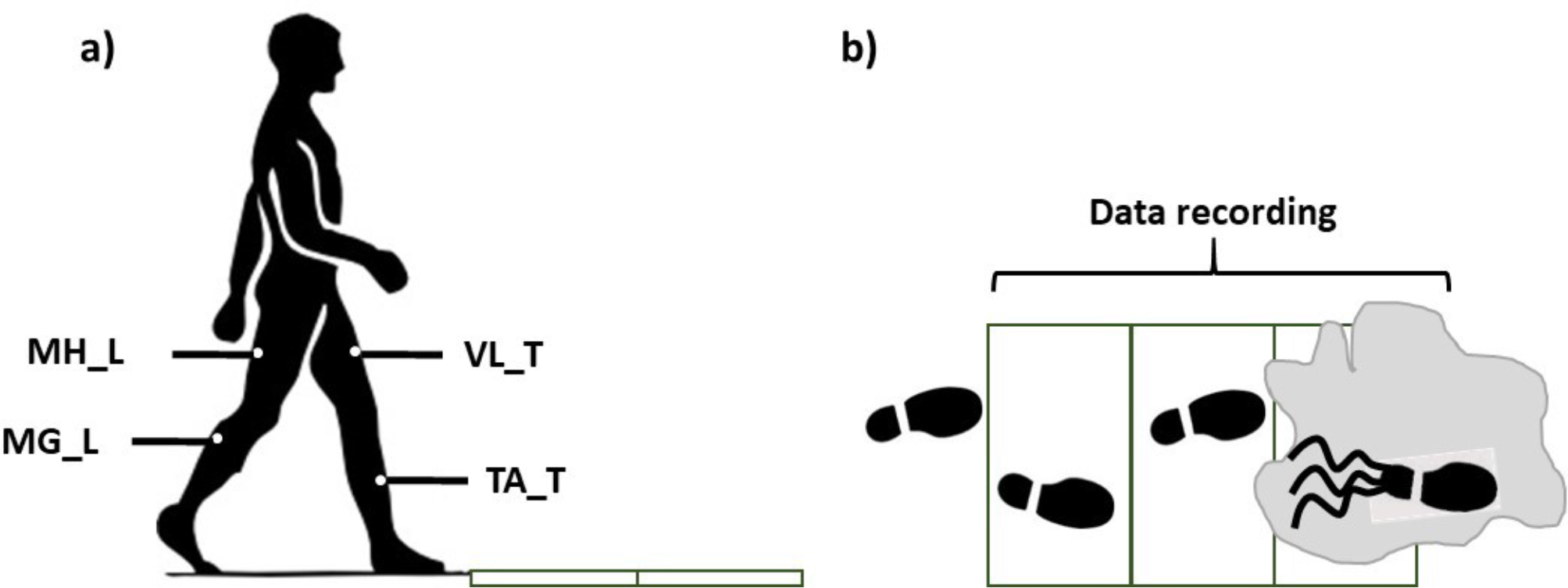
The side view (a) and the top view (b) of the walkway in the final trails (slip). The gray zone indicates the contaminant.

Throughout the walking trials, bilateral EMG signals were recorded at 1080 Hz from medial hamstring (MH), tibialis anterior (TA), vastus lateralis (VL), and medial gastrocnemius (MG) (right/leading/slipping leg (L) and left/trailing/non-slipping leg (T)) (Figure 1). A motion capture system (Vicon 612, Oxford, UK) was utilized to capture heel kinematics at 120 Hz.

### 2.3 Data Analysis

The peak heel speed (PHS) of each subject was used as representative of slip severity using the markers data. Upon recording the walking data, the slip data was used to classify subjects into severe and mild slippers. Persons with a PHS higher than 1.44 m/s were considered severe slippers (T. Lockhart, Woldstad, and Smith 2003) while others were labeled as mild slippers. t-test was used to identify potential inter-group differences in weight, height, and age between mild and severe slippers. Also, a Pearson’s Chi-squared test was performed to examine if there was a significant difference between genders of mild and severe slippers (Table.1).

**Table 1:**
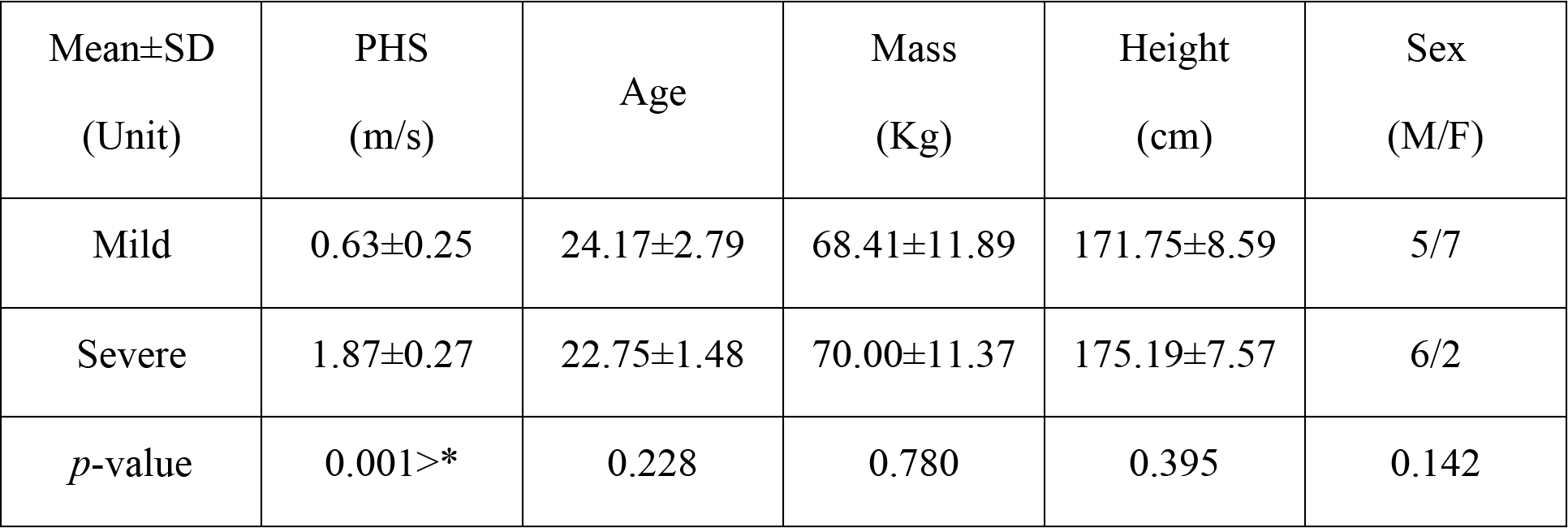
Demographics of different severity groups and the resulting statistical analysis. Please note that Pearson’s Chi squared test was used for Sex while independent *t*-test was used for the others.

EMG signals were processed (demeaned, rectified, filtered) for the walking trial according to previously-described procedures (Nazifi et al. 2017). EMG was then normalized to the maximum activity of each muscle for each subject throughout all of his/her walking and slipping trials. The force plate data was only used to detect the heel strike moment and the gait duration was normalized to 100 points (0 being the first right heel strike, 50 being left heel strike, 100 being the second right heel strike) and an iterative non-negative matrix factorization technique was used to extract walking muscle synergies (Nazifi et al. 2017) and their coefficients from the normalized gait cycle for each subject (Nazifi, Beschorner, and Hur 2017). Prior studies (Nazifi, Beschorner, and Hur 2017;Nazifi et al. 2017) indicated that for walking, four muscle synergies are enough to reconstruct the EMG signals of walking and reach a VAF larger than 95%, hence the number of synergies were fixed to four in this study. Muscle synergies of each severity group were then sorted and re-ordered according to their similarity using correlation coefficients (r) (d’Avella, Saltiel, and Bizzi 2003; Torres-Oviedo and Ting 2010;Nazifi et al. 2017) to have the same synergies (i.e. ones with the highest correlation) in different subjects in the same order (Figure 2). This step was necessary as our method extracted synergies in a random order for each subject. An independent t-test (α=0.05) was used (SPSS v21, IBM, Chicago, IL) to detect the significant differences in muscle synergies between mild and severe slippers. We then used Bonferroni’s 95% confidence interval to examine if the time courses of activation coefficients diverge between the mild and severe slippers.

**Figure 2:**
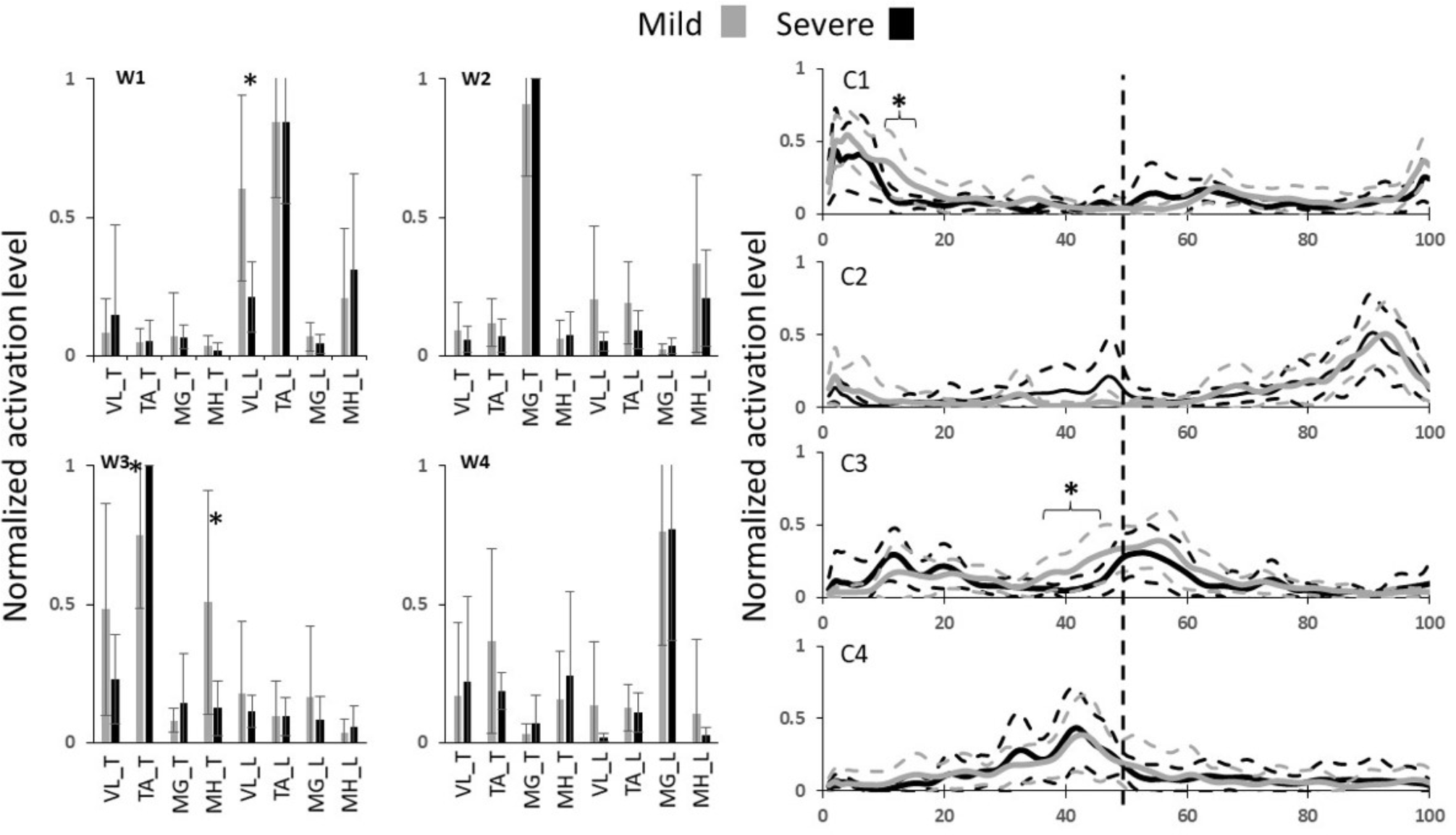
The walking muscle synergies and their activation coefficients for both severity groups. Dashed lines and error bars indicate SD. Asterisks represent significant differences. (0% coincides with leading limb’s heel strike and the trailing limb’s heel strike happens at 50%, i.e. the vertical dashed line).

## 3 Results

From the unexpected slip trial, PHS measurement classified twelve subjects as mild slippers and eight subjects as severe slippers. No significant differences were found in sex, height, mass, and age across severity levels (Table. 1). According to our previous studies, four muscle synergies were enough to account for more than 95% of the EMG variability during walking (Nazifi et al. 2017). Hence, four walking muscle synergies were extracted from each subject (Figure 2). We wish to emphasize that the slipping trial was only performed to classify subjects into potential mild and severe slippers, and the synergy analysis was performed only on walking trials.

Statistical analysis revealed differences in both walking muscle synergies and their activation coefficients (Figure 2). Significant differences contributions of three muscles were found. The different contributions belonged to MH_T, TA_T, and VL_L muscles. MH_T and VL_L muscle had a significantly higher activation in mild slippers, while a higher activation of TA_T was associated with severe slipping (Figure 2, walking muscle synergy 1 and 3, i.e. W1 and W3). The inter-group comparison also found differences in activation of the first and the third walking muscle synergies (Figure 2, C1 and C3). Bonferroni’s 95% confidence interval showed a divergence in the first synergy’s coefficient ‘C1’ between mild and severe slippers from 11th until 15th percent (Figure 2) of the gait cycle (Table 1). A higher activation of C1 in the aforementioned period was associated with mild slips. Also, the activation coefficient of the third muscle synergy, ‘C3’was higher in mild slippers from 37th percent to 45th percent (Figure 2) of the gait cycle according to the same analysis.

## 4 Discussion

The results have indicated that a higher activation of the MH muscle right before the heel strike is associated with less severe slips (Figure 2, MH_T right before the trailing limb’s heel strike at 50%). This point can be seen in both higher contribution of MH_T in the third synergy (W3), and in its higher activation coefficient (C3), right before the trailing limb’s heel strike (which happens around 50%). Hamstring muscle is known to be involved in deceleration of the swing (same as trailing here) leg in right before the heel strike (Basmajian and De Luca 1985; Rose and Gamble 2005; Medved 2000; T. E. Lockhart and Kim 2006), hence, contributing to less slip severity by reducing the heel velocity at the moment of heel strike. According to a prior study from this group (Nazifi, Beschorner, and Hur 2017), a relatively similar sub-task was found to be associated with one of the *slipping* muscle synergies, where subjects with a higher contribution of Hamstring group *during slipping* experienced less severe slips (Nazifi, Beschorner, and Hur 2017). This fact further clarifies the key role that Hamstring group play in fall prevention and slip recovery. Interestingly, as the MH contribution is higher both after a novel slip initiation according to (Nazifi, Beschorner, and Hur 2017; Yang and Pai 2010) and before slip initiation (i.e. in this study, walking trials), it is probable that not only do the hamstring group has a reactive role in fall prevention, but also it may have a proactive role as well.

Moreover, prior studies on walking muscle synergies found one of the synergies be responsible for the ‘load acceptance’ synergy (Nazifi et al. 2017). This synergy prepared the weight to be shifted from the trailing limb to the leading limb. Despite the different scope of the studies, a comparable pattern was found in W1 in this study and is considered to be associated with the load acceptance. Statistical analysis of this synergy shows that a higher activation of the VL muscle right after the heel strike is associated with less severity in slips. This conclusion was made upon the observation of a higher contribution of VL_L in W1 along with a higher activation in C1, right after the leading limb’s heel strike (which happens around 0%, Figure 2). Considering the role of VL in load acceptance, this conclusion stays consistent with existing studies claiming that a late activation of the VL may reduce the forward velocity of the center of mass relative to the base of support, resulting in less stability (Cham and Redfern 2001; Chambers and Cham 2007). In other words, mild slippers had a higher activation on their load acceptor muscle (VL_L) shortly after leading limb’s heel contact enabling them to transfer their weight with more support.

The third muscle synergy suggests a toe lift behavior. Based on the contribution of each muscle it is likely that this synergy contributes to elevation of the toes right before the heel strike, probably to avoid tripping or foot drop. However, there was an association between higher activation of the TA muscle before the heel strike and high slip severity. TA_T muscle had a higher contribution in the third muscle synergy (W3). It was previously shown that severe slippers’ high activation of TA right before their heel strike increases their foot-floor-angle significantly (FFA) and was found to be associated with their severe slips (Nazifi, Beschorner, and Hur 2017). An excessive dorsiflexion and FFA right before the heel strike also challenges achievement of flat-foot and recovery (Chambers and Cham 2007). It is also known that a reduced FFA (i.e. flat-foot walking) is a strategy used by individuals to increase dynamic stability of the gait (Gao and Abeysekera 2004; Strandberg and Lanshammar 1981; Marigold and Patla 2002; Bhatt, Wening, and Pai 2006). Finding of an excessive contribution from TA muscle in walking muscle synergies of severe slippers verifies our findings about a correlation between FFA and propensity to falls while normal walking.

## 5 Conclusion

This study examined the walking muscle synergies and their differences for different slip severities. We found significant differences in the walking muscle synergies of mild and severe slippers. This study provides a basis for a potential diagnosis method to identify the vulnerable population and people with high risk of fall based on solely their walking pattern and improves their safety and consequently, quality of life. Such a diagnosis method will be valuable as it does not require an actual slip trial once a predictive model is developed. There were a few limitations to our study. Our study was only performed on the young adults and can be significantly improved by including older populations. Also, as falls impose more detrimental consequences on the older adults, our future studies would test the differences between mild and severe slippers in older populations. Moreover, this study only targeted unexpected slips. A potential different can be present between the response to unexpected and expected slips that can be addressed in future. Also, only eight major muscles (i.e. four muscles per limb) were studied. Future studies can resolve this limitation by studying more muscles that may contribute to human gait. Future studies also will develop and study the effectiveness of a predictive model in identifying severe slippers. Another limitation of this study is the limited number of subjects. A future study is needed to confirm the findings in a larger group. Also, despite that our statistical analysis did not find the slip severity to be gender-related, since prior studies have shown gender-related discrepancies in walking patterns (Cho, Park, and Kwon 2004), gender’s association with slip severity will be studied in a larger data set. Lastly, this study has not controlled for footedness of the subjects and it can be improved by controlling the footedness of each subject upon heel strike on the slippery surface.

## Acknowledgement

We thank Dr. Cham for providing the experimental data for the analyses.

## Abbreviations

CNS: Central Nervous System
SD: Standard Deviation
PHS: Peak Heel Speed (during a slip)
PVC: Polyvinyl Chloride
L: Leading
T: Trailing
MH: medial hamstring
TA: tibialis anterior
VL: vastus lateralis
MG: medial gastrocnemius
VAF: Variance Accounted For
FFA: Foot-floor Angle

